# Extensive transcriptional differentiation and specialization of a parasite across its host’s metamorphosis

**DOI:** 10.1101/2024.07.16.603694

**Authors:** James G. DuBose, Jacobus C. de Roode

## Abstract

Foundational theory on life cycle evolution suggests that given genetic independence, the phenotypes presented by different life stages will diverge more when they occupy more dissimilar niches. When divergence between stages is significant and punctual, we typically consider the life cycle complex. In parasites, the delineation between simple and complex life cycles is usually made between those that occupy single and multiple host species. However, many parasites can experience significant niche shifts in a single host. To explore the potential for a host’s metamorphosis to shape divergence between stages across its parasite’s life cycle, we quantified the transcriptional differentiation and specialization that the protozoan parasite *Ophryocystis elektroscirrha* exhibits across the metamorphosis of its host the monarch butterfly. We found evidence that *O. elektroscirrha* differentiates in concordance with the ecological turnover imposed by monarch transitions to different stages, and that patterns of transcriptional decoupling across the *O. elektroscirrha* exceeded even that of its host. However, because of its greater gene content, the monarch exhibited greater total transcriptional turnover than its parasite. These findings highlight that synthesis of evolutionary theory pertaining to free-living and parasitic life cycles could be facilitated by more nuanced and continuous descriptions of life cycle complexity.

## Introduction

The life cycles of many organisms require progression through series of phenotypic transitions that facilitate their activities in different ecological niches. This phenotypic turnover is best exemplified by life cycles that involve complex and abrupt changes, such as the metamorphosis of tadpoles into frogs or the transmission between distinct host species of parasites. Deeply rooted divergence between lineages has often occurred in association with principal life cycle alterations (Wheeler et al. 2001; Reiss 2002). Therefore, an understanding of life cycle evolution has the potential to inform a more general understanding of patterns of biological diversification.

The fundamental goal of studying life cycle evolution is to understand how and why the phenotypes presented by an organism differentiate as they undergo ontogenetic niche shifts (Werner and Gilliam 1984). Theory suggests that if the traits presented by different stages share a genetic basis, phenotypic divergence between stages will be relatively constrained because variation that is neutral or beneficial in one niche may be detrimental in another (Ebenman 1992). However, if the traits expressed by different stages become genetically decoupled, different stages may evolve independently, thus facilitating their differentiation (Ebenman 1992).

Genetic (and consequently evolutionary) decoupling of at least some traits expressed by different life stages is empirically well supported (Fellous and Lazzaro 2011; Aguirre et al. 2014; Goedert and Calsbeek 2019; Schott et al. 2022). Underlying this genetic independence and phenotypic differentiation is transcriptional dissimilarity between life stages, just as transcriptional dissimilarity generates differentiation between cell types (Fellous and Lazzaro 2011; Herrig et al. 2021; Collet et al. 2023; DuBose and de Roode 2024). Therefore, a primary driver of differentiation is the specialization of certain genes’ expression to a given stage, as genes that are expressed broadly at consistent levels across an organism’s life cycle contribute little to transcriptional dissimilarity between stages. Increased temporal specificity in patterns of gene expression have been linked to gene duplication and increased rates of molecular evolution, which shows general consistency theory on life cycle evolution and our understanding of how cell and tissue differentiation evolves (Williams et al. 2023; DuBose and de Roode 2024).

Although we are beginning to understand the genetic and transcriptional basis of trait decoupling across ontogeny, an inconsistency remains between theory describing life cycle evolution and how life cycle complexity is typically evaluated, particularly in parasites. From a zoological perspective, which much theory on life cycle evolution is based on, life cycles are considered complex if they progress through two or more discrete stages (Istock 1967; Moran 1994). Similarly, parasites are considered to have complex life cycles if they progress through two or more host species, and simple if they occupy a single host species (Combes 2001; Auld and Tinsley 2015). These perspectives are conceptually similar in that they consider abrupt ontogenetic niche shifting indicative of greater life cycle complexity (Benesh 2016). However, many parasites can experience significant niche shifts in a single host, particularly those that move between different tissues (Haldane 1932) or, as we explore here, occupy hosts that undergo a metamorphosis.

More generally, theory suggests that the greater the dissimilarity between niches occupied by different stages, the more their phenotypes (given genetic independence) will diverge (Ebenman 1992). Here we apply this concept towards describing the transcriptional dynamics of *Ophryocystis elektroscirrha*, a protozoan parasite that infects the monarch butterfly *Danaus plexippus* as its single host (McLaughlin and Myers 1970). We found evidence that *O. elektroscirrha* undergoes significant functional differentiation throughout the monarch metamorphosis, and that patterns of transcriptional dissimilarity across the *O. elektroscirrha* life cycle correspond with patterns of transcriptional dissimilarity between monarch stages. Given that the monarch life cycle is considered complex while that of the parasite is considered simple, these findings suggest we need to reevaluate how we describe life cycle complexity.

## Methods

### Study system

*O. elektroscirrha* is an apicomplexan parasite that completes its development in the monarch butterfly (Figure 1). *O. elektroscirrha* forms dormant spores (oocysts) on the adult monarch abdomen, which are passively transferred to milkweed foliage and eggs when adult butterflies touch milkweed or oviposit (de Roode et al. 2009; Majewska et al. 2019). Infection starts when monarch larvae ingest said spores. Spores lyse in the larval gut and then penetrate the gut wall to infect hypodermal cells. As monarch larvae transition to pupae, parasites continue to replicate and rupture the hypodermal cells. Early in the monarch pupal stage, parasites replicate once more before maturing in the hypoderm and hemolymph during the middle of the pupal stage. Later during the monarch pupal development, parasites will congregate and pair to form zygotes. Just prior to monarch eclosion, parasites will undergo meiosis, where sporulation involves 3 nuclear divisions, resulting in spores with 8 parasites enclosed. When the monarch ecloses and the adult butterfly emerges, parasites spores are concentrated on their abdomens (Leong et al. 1992), from which they will be transmitted to the next generation (McLaughlin and Myers 1970).

**Figure 1.**
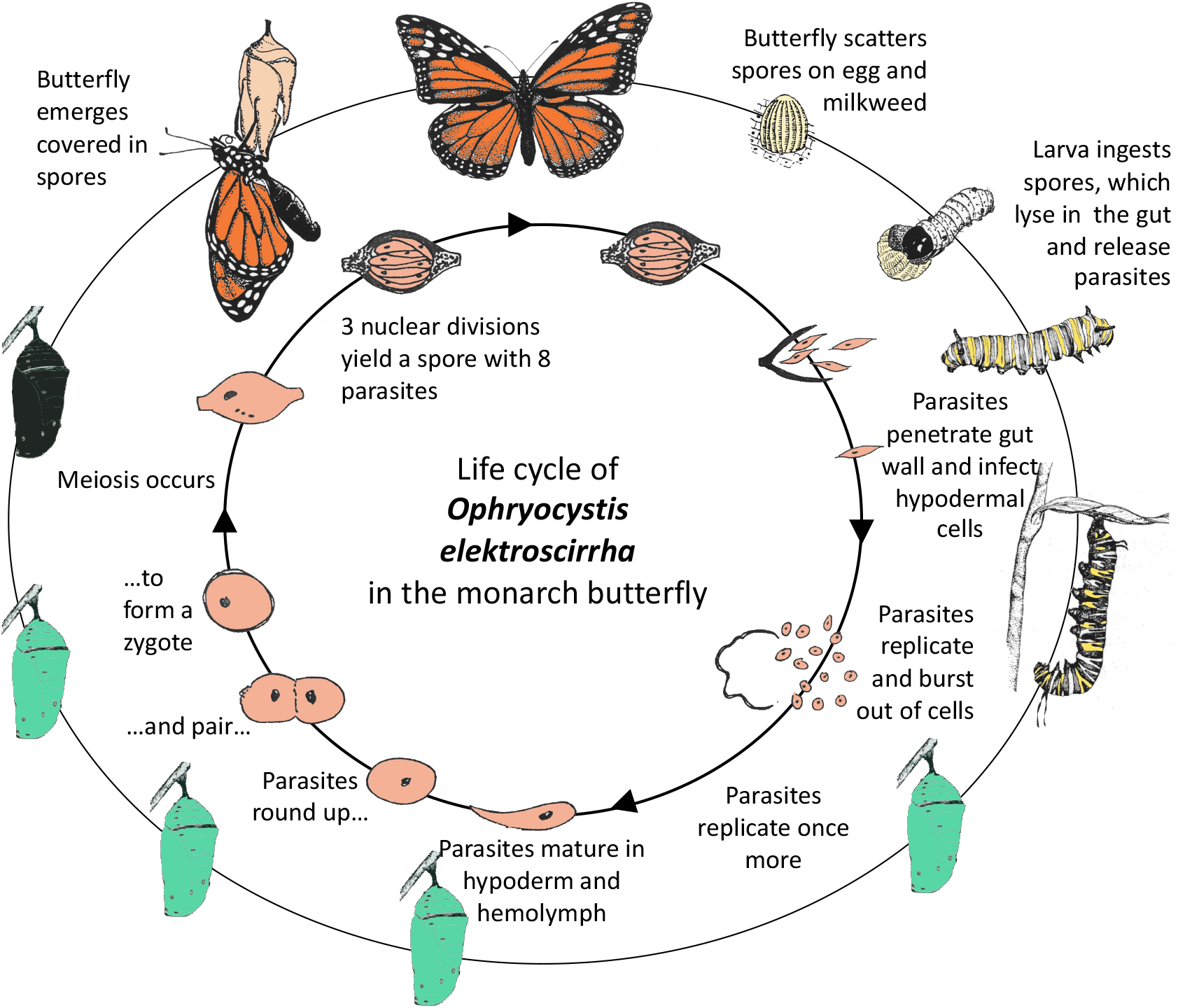
An illustration of the *O. elektroscirrha* life cycle. Image depicts the timing of *O. elektroscirrha* growth and reproduction events relative to the monarch metamorphic transitions.

### Study design and implementation

To quantify the functional differentiation of *O. elektroscirrha* across its life cycle, we sequenced mRNA from extracted from infected monarch third instar larvae, fifth instar larvae, early pupae (one day post-pupation), late pupae (six to eight days post pupation), and adults (several hours after eclosion). We used the protocols described in (DuBose and de Roode 2024) for all monarch rearing and mating. Briefly, the *D. plexippus* individuals we used in this study were F2 descents from a parental generation that was wild caught near St. Marks, Florida, U.S.A. (30°09′33″N 84°12′26″W). The milkweeds that monarch larvae eat contain toxic compounds, and high concentrations of these compounds to negatively impact *O. elektroscirrha* proliferation (Hoogshagen et al. 2024). Therefore, we reared *D. plexippus* caterpillars on *Asclepias incarnata* (less toxic) and *Asclepias curassavica* (more toxic) to ensure our findings were robust to a potentially important source of environmental variation that could influence *O. elektroscirrha* development. To administer infections, we placed 100 *O. elektroscirrha* spores (isolate 23P4) on leaf discs from the corresponding treatment plant and fed them to second instar monarch larvae, following existing protocols (de Roode et al. 2006). We collected four to five replicates for each stage on each plant (Appendix, Table S1). Because we did not find a significant effect of larval diet on *O. elektroscirrha* developmental dynamics (Appendix, Section 3.1), we did not consider plant treatments in subsequent analyses. Therefore, our final sampling consisted of eight to ten replicates per life stage.

### Sample collection and mRNA sequencing

We used the protocols established in (DuBose and de Roode 2024) for all monarch homogenate collection, RNA extraction, and RNA sequencing. Briefly, we snap froze all monarch individuals in liquid nitrogen and stored them at 80°C prior to tissue homogenization and RNA extraction. Because *O. elektroscirrha* occupies several different monarch tissues, we extracted RNA from whole body monarch homogenate using a Promega SV Total Isolation System. We made several alterations to the manufacturer’s suggested protocol to obtain higher quality RNA extracts. Briefly, we doubled the volume of the recommended RNA lysis buffer, increased the relative centrifugal force in all centrifugation steps, and included additional centrifugation to better clear organic contaminants and improve final extract purity. All quantifications of RNA extract purity can be found in Appendix, Table S2. After all extractions were completed, we sent purified RNA to Novogene (Sacramento, CA) for library preparation and sequencing. Novogene used an Agilent 5400 Fragment Analyzer System to assess RNA extract purity and integrity (minimum RIN = 7.9). Sequencing libraries were then prepared via poly-A tail selection and sequenced on a NovaSeq 6000 sequencing system using a 150bp paired-end approach, ensuring that a minimum of 20 million reads were obtained for each sample.

### Sequence processing and transcript quantification

Novogene performed initial quality control of raw sequences, where sequences with a phred score of less than or equal to 5 in 50% of the read, sequences that consisted of 10% or higher ambiguous base calls, and adapter sequences were discarded. We then performed additional quality assessment using FASTQC (Andrews 2010), which showed that the median phred score did not drop below 30 at any sequence position for any sample.

To quantify transcript abundances, we first used STAR (v. 2.7.11a) (Dobin et al. 2013) to align reads to the coding sequences of the *D. plexippus* genome (v.Dpv3, GenBank Assembly = GCA_000235995.2) and the *O. elektroscirrha* genome (GenBank Assembly = GCA_030141585.1) (Mongue et al. 2023). Reads that aligned to both genomes were considered ambiguous and discarded from further analyses and reads aligned to only one reference genome were split into separate files. Overall, we found an average of 538214.35 *O. elektroscirrha* reads/sample (SD = 340284.89, range = [145062, 1977293], see Appendix, Section 2.2 for a full summary), which given its small gene content and low cell number relative to host cells, represents an impressive recovery of *O. elektroscirrha* transcripts. We then quantified transcript abundances using kallisto (v.0.46.2) (Bray et al. 2016). We performed downstream analyses using log transformed transcript per million normalized read counts (automatically generated by kallisto) to minimize biases due to the gene expression data distributions, unequal gene lengths, and varying library sizes (Wagner et al. 2012; Abrams et al. 2019).

### Analyses of functional differentiation and transcriptional dissimilarity

To examine *O. elektroscirrha’s* potential functional differentiation between stages, we used GOATOOLS (v. 1.4.12) (Klopfenstein et al. 2018) to assign cellular component and biological process Gene Ontology annotations (The Gene Ontology Consortium et al. 2023) to transcripts. We then calculated *O. elektroscirrha’s* relative investment in each category by summing the expression values for each gene that shared a Gene Ontology annotation. We performed this separately for cellular components and biological processes. We then performed a principal component analysis using *PCA* function from the *scikit-learn decomposition* module (Pedregosa et al. 2011) to visualize dissimilarity in functional investment between *O. elektroscirrha* in different *D. plexippus* stages, as well as identify the cellular components and biological processes that best explained patterns of differentiation. We then statistically tested for significant differentiation in *O. elektroscirrha* cellular components and biological processes across *D. plexippus* life stages by using the *adonis2* function from the *vegan* R package (v.2.6-4) (Oksanen et al. 2022) to perform a permutational multivariate analysis of variance (PERMANOVA) with 999 permutations.

To test that patterns of transcriptional dissimilarity across the *O. elektroscirrha* life cycle corresponded to its host’s transcriptional changes, we first calculated dissimilarity matrices based on Pearson correlation distances 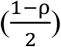 for *O. elektroscirrha* and its host. We then performed a Mantel test using the *mantel* function in the *vegan* R package (Oksanen et al. 2022) to calculate the correlation between the two distance matrices and test for its statical significance. To quantify how much variation in patterns of *O. elektroscirrha* transcriptional dissimilarity could be explained by the patterns of transcription dissimilarity in its host, we performed a redundancy analysis using the *rda* function from the *vegan* R package (Oksanen et al. 2022).

Because the temporal specificity of a gene’s expression is an important aspect of differentiation, we quantified the stage-specificity across all *O. elektroscirrha* and *D. plexippus* gene expression profiles. First, we took the average expression for each gene. We used the tissue-specificity index τ to quantify stage-specificity, which ranges from 0 (equal expression across stages) to 1 (expression in a single stage): 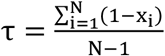, where *N* is the number of stages (for our purposes) and *x*_*i*_ is the expression level normalized to the maximum expression value across stages. We performed a Kolmogorov–Smirnov test using the *ks*.*test* R function (R Core Team 2022) to compare the distributions of τ between *O. elektroscirrha* and its host.

Finally, we compared the extent of transcriptional turnover across the *O. elektroscirrha* life cycle to that of its host. First, we calculated the average Pearson correlation distance between samples in adjacent stages, which represents the extent that transcription in one stage correlates with transcription in the other (higher values indicate weaker correlation). However, this approach is not sensitive to the number of genes possessed by each organism, and we wanted to evaluate the concept that the monarch host could exhibit greater transcriptional turnover due to its higher gene content. Therefore, we performed the same analysis using Manhattan distances as described in (DuBose and de Roode 2024), which is more representative of total transcriptional turnover.

## Results

### O. elektroscirrha undergoes significant functional differentiation throughout the D. plexippus metamorphosis

We found that *O. elektroscirrha* exhibited significant functional differentiation, both in terms of cellular components (F = 65.93, p ≤ 0.001) and biological processes (F = 59.71, p ≤ 0.001), between different *D. plexippus* life stages (Figure 2). During the early pupal stage, *O. elektroscirrha* functional differentiation was associated with increased investment in genes involved in nucleolus, small subunit processome, ribosomal and nuclear activities. These investments are consistent with the rapid growth and the vast multi-nucleation associated with micronuclear schizogony (a form of asexual reproduction) during this stage (McLaughlin and Myers 1970) (Figures 1, Figure 2A). Consistent with this period of proliferation and multi-nucleation, *O. elektroscirrha* differentiation was also associated with investment in genes involved in translation, RNA binding, and nucleic acid binding (Figures 1, Figure 2B). During the early pupal stage, *O. elektroscirrha* also increased investment in genes involved in proteasome production and activities. However, increased investment in proteolysis was not seen until the late pupal stage, which could indicate preemptive investment in the structures needed for physiological adjustment during the late pupal stage. During the later pupal stage *O. elektroscirrha* differentiation was associated with investment in genes involved in membrane dynamics, as well as ATP binding and protein binding (Figure 1, Figure 2B). These investments are consistent with the energetic demands of motility via cytoplasmic constriction at this stage, as well as the eventual cell-cell recognition and membrane fusion involved in zygote formation later during this period (McLaughlin and Myers 1970) (Figure 1, Figure 2B).

**Figure 2.**
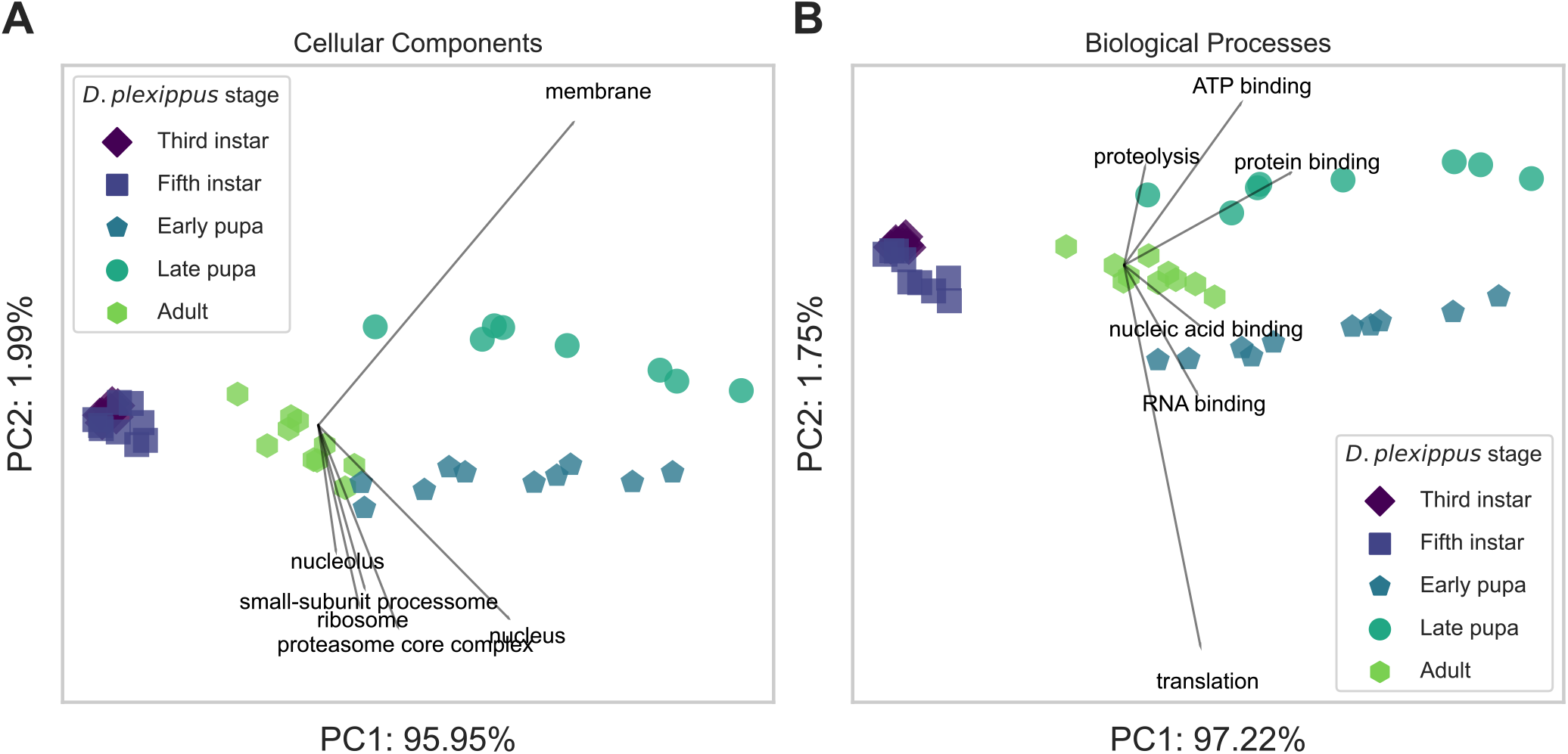
*O. elektroscirrha* undergoes functional differentiation across the *D. plexippus* life cycle. Principal component plots show the dissimilarity in A) cellular components and B) biological processes that *O. elektroscirrha* invests in throughout the *D. plexippus* life cycle. Each point represents an individual sample, and closer points indicate the samples had more similar functional profiles. Labeled vectors indicate the six functions that explained the most variation in patterns of functional dissimilarity. Vectors were re-scaled to provide better visualization, but their relative lengths were preserved. Axis labels indicate the principal component rank and the proportion of variance it explained.

### Patterns of transcriptional variation across the D. plexippus metamorphosis predict the patterns of transcriptional variation across the O. elektroscirrha life cycle

Theory suggests that differentiation between life stages facilitates an organism’s activities and specialization in different ecological niches across its life. Therefore, we tested that the patterns of differentiation between *O. elektroscirrha* stages would reflect the patterns of dissimilarity in the niches they occupy, which evaluated as transcriptional dissimilarity between its host’s stages. Consistent with predictions, we found that patterns of transcriptional dissimilarity across the *O. elektroscirrha* life cycle were highly correlated with patterns of transcriptional dissimilarity across the *D. plexippus* life cycle (r_M_ = 0.8132, p < 0.001) (Figure 3). Furthermore, patterns of transcriptional dissimilarity across the *D. plexippus* life cycle explained 87.15% of the variation in transcriptional dissimilarity across the *O. elektroscirrha* life cycle.

**Figure 3.**
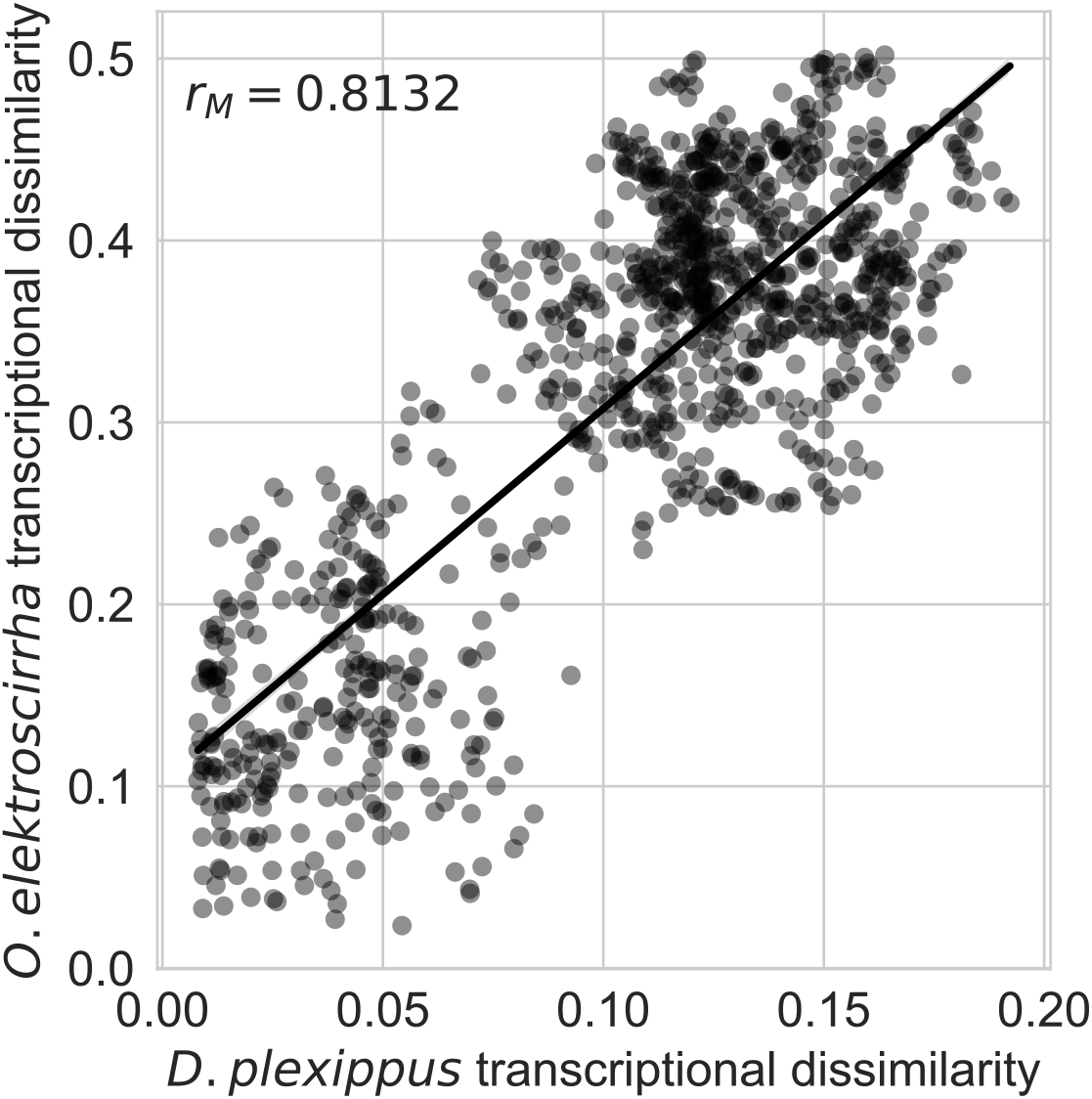
Transcriptional dissimilarity between *D. plexippus* life stages predicts patterns of transcriptional dissimilarity across the *O. elektroscirrha* life cycle. The x-axis indicates the Pearson distance between each pair of *D. plexippus* samples, and the y-axis indicates the Pearson distance between their corresponding *O. elektroscirrha* samples. The regression line was plotted to aid in visualization of the correlation, not for statistical testing (as pairwise comparisons are not independent).

### O. elektroscirrha shows greater transcriptional specialization and differentiation than D. plexippus, but less total transcriptional turnover

The fundamental driver of differentiation between life stages is temporal specificity in patterns of gene expression: genes with greater temporal specificity contribute more to differentiation because their effects are limited to the time in which they are expressed (Haldane 1932). Therefore, we quantified the distribution of stage specificity across *O. elektroscirrha* genes and compared it to that of its host. Surprisingly, we found that *O. elektroscirrha* genes showed far greater stage specificity than monarch genes (D = 0.58, p < 2.2×10^−16^) (Figure 4A). Consistent with more stage-specific patterns of gene expression, the correlation in gene expression between subsequent stages was lower in *O. elektroscirrha* across each transition (Figure 4B). However, we reasoned that because *D. plexippus* has more genes than *O. elektroscirrha*, relatively subtle differences in monarch gene expression could cumulate to produce greater total transcriptional turnover. Consistent with this prediction, we found that when transcriptional dissimilarity was evaluated using Manhattan distances, *D. plexippus* showed greater differences in gene expression between each set of subsequent stages (Figure 4C).

**Figure 4.**
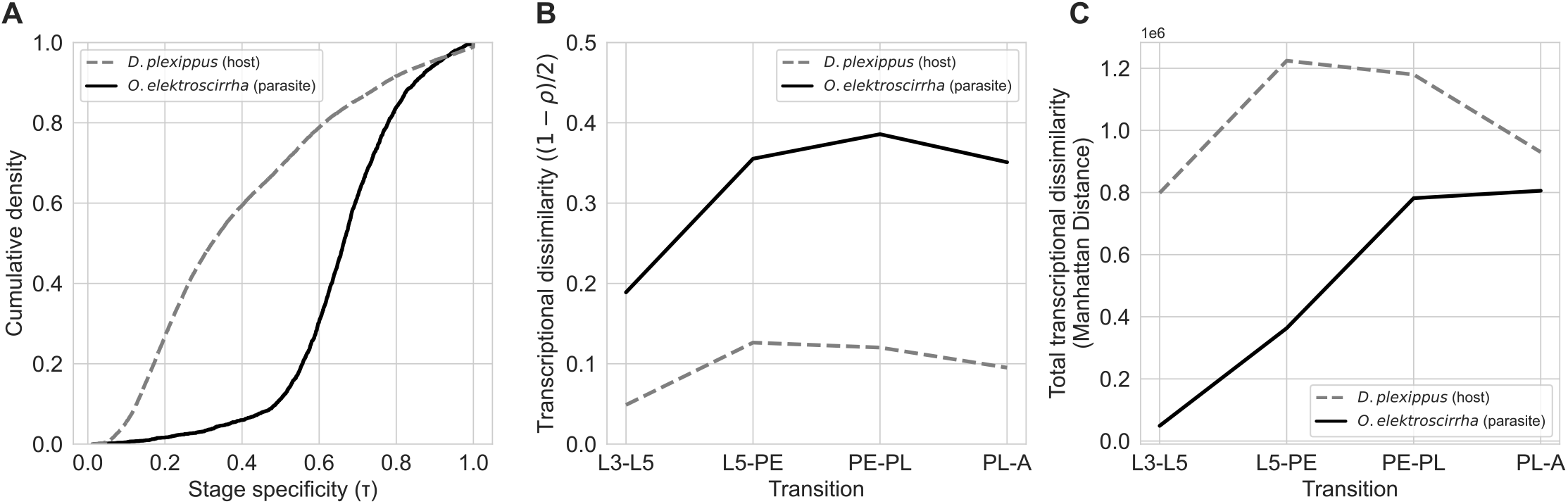
The extent of transcriptional decoupling is greater across the *O. elektroscirrha* life cycle, but total transcriptional turnover is greater across the *D. plexippus* life cycle. A) The empirical cumulative density function of stage specificity values for all *O. elektroscirrha* (solid) and *D. plexippus* (dashed) genes. B) The average Pearson distance or C) Manhattan distance between each *D. plexippus* transition for *O. elektroscirrha* (solid) and *D. plexippus* (dashed) gene expression profiles. The x-axis is organized into sequential transitions, where L3 = third instar larvae, L5 = fifth instar larvae, PE = early pupae, PL = late pupae, and A = adults.

## Discussion

Life cycle complexity is generally marked by abrupt ontogenetic niche shifts across an organisms’ life (Benesh 2016). From the parasitological perspective, this niche shift has been predominately associated with transitioning to distinct host species throughout development (Combes 2001; Auld and Tinsley 2015). However, here we explored the potential for a parasite whose life cycle would typically be considered simple to undergo abrupt ontogenetic niche shifts throughout its host’s complex (metamorphic) life cycles. By examining temporal patterns of gene expression in the monarch butterfly and its parasite *O. elektroscirrha*, we found evidence that *O. elektroscirrha* undergoes significant functional differentiation and specialization as its host transitions to different life stages (Figures 1 and 2). In support of *O. elektroscirrha’s* differentiation occurring in concordance with an abrupt ontogenetic niche shifts, we found that transcriptional dissimilarity between host stages predicted patterns of *O. elektroscirrha* transcriptional dissimilarity across its life cycle (Figure 3). Finally, we found that *O. elektroscirrha’s* genes showed more stage-specific expression profiles than its host’s genes, which translated into weaker transcriptional correlation between stages across the *O. elektroscirrha* life cycle (Figure 4A and B). However, because of its greater gene content, the total transcriptional turnover exhibited by *D. plexippus* exceeded that of its parasite (Figure 4C).

The findings that pattern of *O. elektroscirrha* gene expression showed increased stage specificity and weaker correlations between stages than that of its host indicates its potential for greater trait decoupling between stages. However, we do not interpret this as evidence for *O. elektroscirrha’s* greater life cycle complexity, as monarchs still showed more total transcriptional turnover (which would likely be even higher if we analyzed different monarch tissue types separately). Rather, these findings suggest that the widely employed binary classification of life cycles as “simple” or “complex” may be insufficient to describe life cycle evolution. A more general interpretation would be that *O. elektroscirrha* may show greater trait decoupling between stages, but its reduced gene content limits the total transcriptional turnover across its life cycle.

Conversely, monarchs showed comparatively less decoupling between stages, but exhibited greater total transcriptional turnover due to their increased organismal complexity (more genes and cell types). The fundamental link between the two is that both organisms modulate their transcription as their ecological niches change, which is consistent with Haldane’s earlier thoughts that a parasite that occupies multiple different host tissues might express different sets of genes to suit its activities in different tissue types (Haldane 1932). Therefore, a more fundamental understanding of life cycle evolution could come from a general understanding of how organisms evolve specialized expression patterns of gene expression (Haldane 1932; Makova and Li 2003; Li et al. 2005; DuBose and de Roode 2024). Furthermore, these findings raise more general questions about the role of genomic complexity in limiting or promoting transcriptional specialization to specific life stages or time points.

These findings also exemplify several other issues with the binary classification of life cycles. From the zoological perspective, the subjectivity of defining discrete stages is a recognized issue (Moran 1994; Bishop et al. 2006; DuBose and de Roode 2024). From the parasitological perspective, transitioning from one host species to another may certainly represent a significant ecological shift. However, parasites that occupy a single host (whose life cycle would be considered simple) can also experience a significant niche shift as they migrate to different tissues (Haldane 1932) or as we describe here, as their host undergoes a significant metamorphosis. It is important to note however, that we do not suggest classifying parasitic life cycles based on the number of hosts they occupy has no utility. The number of hosts a parasite occupies certainly has ecological and epidemiological consequences. Rather, integration of evolutionary theory pertaining to free-living and parasitic life cycles could benefit by considering life cycle complexity on a continuum.

## Author Contributions

J.G.D and J.C.dR designed and performed research. J.G.D analyzed the data. J.G.D and J.C.dR wrote the paper.

## Data and code availability

All sequences and count matrices generated for this project have been deposited in the NCBI GEO database and will be made available upon publication. All code written for data analysis can be accessed at https://github.com/gabe-dubose/oeld.

