## Appendix for "Extensive transcriptional differentiation and specialization of a parasite across its host’s metamorphosis"

### Appendix for:
### *The extensive transcriptional differentiation and specialization of a parasite across its host's metamorphosis*

James G. DuBose[1,*] and Jacobus de Roode[1]

[1] *Department of Biology, Emory University*

### 1 Extended Methods

#### 1.1 Sample metadata

Table S1: Sample metadata

| Sample ID | Larval plant host | Infection status | Monarch Stage |
|---|---|---|---|
| mtstp3ci10 | curassavica | infected | third-instar |
| mtstp3ci11 | curassavica | infected | third-instar |
| mtstp3ci12 | curassavica | infected | third-instar |
| mtstp3ci13 | curassavica | infected | third-instar |
| mtstp3ci9 | curassavica | infected | third-instar |
| mtstp3ii89 | incarnata | infected | third-instar |
| mtstp3ii90 | incarnata | infected | third-instar |
| mtstp3ii92 | incarnata | infected | third-instar |
| mtstp3ii94 | incarnata | infected | third-instar |
| mtstp5ci25 | curassavica | infected | fifth-instar |
| mtstp5ci26 | curassavica | infected | fifth-instar |
| mtstp5ci27 | curassavica | infected | fifth-instar |
| mtstp5ci28 | curassavica | infected | fifth-instar |
| mtstp5ii105 | incarnata | infected | fifth-instar |
| mtstp5ii107 | incarnata | infected | fifth-instar |
| mtstp5ii108 | incarnata | infected | fifth-instar |
| mtstp5ii109 | incarnata | infected | fifth-instar |
| mtstp5ii112 | incarnata | infected | fifth-instar |
| mtstpAci73 | curassavica | infected | adult |
| mtstpAci74 | curassavica | infected | adult |
| mtstpAci75 | curassavica | infected | adult |
| mtstpAci76 | curassavica | infected | adult |
| mtstpAci77 | curassavica | infected | adult |
| mtstpAii153 | incarnata | infected | adult |
| mtstpAii154 | incarnata | infected | adult |
| mtstpAii155 | incarnata | infected | adult |
| mtstpAii156 | incarnata | infected | adult |
| mtstpAii157 | incarnata | infected | adult |
| mtstpEci41 | curassavica | infected | early-pupa |
| mtstpEci42 | curassavica | infected | early-pupa |
| mtstpEci43 | curassavica | infected | early-pupa |
| mtstpEci44 | curassavica | infected | early-pupa |
| mtstpEci45 | curassavica | infected | early-pupa |
| mtstpEii121 | incarnata | infected | early-pupa |
| mtstpEii122 | incarnata | infected | early-pupa |
| mtstpEii123 | incarnata | infected | early-pupa |
| mtstpEii124 | incarnata | infected | early-pupa |
| mtstpEii125 | incarnata | infected | early-pupa |
| mtstpLci57 | curassavica | infected | late-pupa |
| mtstpLci58 | curassavica | infected | late-pupa |
| mtstpLci60 | curassavica | infected | late-pupa |
| mtstpLci64 | curassavica | infected | late-pupa |
| mtstpLii137 | incarnata | infected | late-pupa |
| mtstpLii138 | incarnata | infected | late-pupa |
| mtstpLii140 | incarnata | infected | late-pupa |
| mtstpLii141 | incarnata | infected | late-pupa |

### 2 Methodological Summaries

#### 2.1 RNA quality report

Table S2: RNA extract quality control report.

| Sample Name | Concentration (ng/ul) | Volume (ul) | Total amount (ug) | RIN |
|---|---|---|---|---|
| mtstp3ci10 | 96.67 | 92 | 8.89 | 9.5 |
| mtstp3ci11 | 288.28 | 92 | 26.52 | 9.6 |
| mtstp3ci12 | 112.59 | 94 | 10.58 | 9.8 |
| mtstp3ci13 | 191.82 | 94 | 18.03 | 9.8 |
| mtstp3ci9 | 347.62 | 92 | 31.98 | 9.7 |
| mtstp3ii89 | 300.2 | 97 | 29.12 | 9.6 |
| mtstp3ii90 | 170.44 | 91 | 15.51 | 9.8 |
| mtstp3ii92 | 231.65 | 91 | 21.08 | 9.7 |
| mtstp3ii94 | 157.16 | 92 | 14.46 | 9.6 |
| mtstp5ci25 | 240.76 | 91 | 21.91 | 9.8 |
| mtstp5ci26 | 342.08 | 91 | 31.13 | 9.7 |
| mtstp5ci27 | 106.47 | 92 | 9.79 | 9.7 |
| mtstp5ci28 | 286.33 | 91 | 26.06 | 9.8 |
| mtstp5ii105 | 247.12 | 89 | 21.99 | 9.8 |
| mtstp5ii107 | 105.85 | 91 | 9.63 | 9.8 |
| mtstp5ii108 | 488.18 | 94 | 45.89 | 9.7 |
| mtstp5ii109 | 47.89 | 92 | 4.41 | 9.4 |
| mtstp5ii112 | 199.79 | 89 | 17.78 | 9.8 |
| mtstpAci73 | 194.49 | 92 | 17.89 | 9.6 |
| mtstpAci74 | 105.01 | 92 | 9.66 | 9.4 |
| mtstpAci75 | 37.05 | 89 | 3.3 | 9.2 |
| mtstpAci76 | 191.76 | 91 | 17.45 | 9.5 |
| mtstpAci77 | 79.2 | 92 | 7.29 | 9.2 |
| mtstpAii153 | 92.54 | 92 | 8.51 | 9.5 |
| mtstpAii154 | 113.15 | 92 | 10.41 | 9.5 |
| mtstpAii155 | 123.51 | 94 | 11.61 | 9.5 |
| mtstpAii156 | 34.04 | 89 | 3.03 | 8.2 |
| mtstpAii157 | 66.15 | 93 | 6.15 | 9.2 |
| mtstpEci41 | 146.55 | 91 | 13.34 | 8.7 |
| mtstpEci42 | 485.48 | 92 | 44.66 | 8.4 |
| mtstpEci43 | 343.02 | 92 | 31.56 | 7.8 |
| mtstpEci44 | 219.26 | 91 | 19.95 | 9.2 |
| mtstpEci45 | 233.62 | 88 | 20.56 | 8.6 |
| mtstpEii121 | 349.96 | 91 | 31.85 | 8.1 |
| mtstpEii122 | 169.85 | 94 | 15.97 | 9.2 |
| mtstpEii123 | 288.9 | 91 | 26.29 | 8.5 |
| mtstpEii124 | 372.33 | 91 | 33.88 | 8.1 |
| mtstpEii125 | 103.25 | 89 | 9.19 | 8.6 |
| mtstpLci57 | 374.64 | 93 | 34.84 | 8.6 |
| mtstpLci58 | 270.8 | 90 | 24.37 | 9.8 |
| mtstpLci60 | 139.83 | 93 | 13 | 8.7 |
| mtstpLci64 | 281 | 92 | 25.85 | 8.8 |
| mtstpLii137 | 169.53 | 91 | 15.43 | 8.5 |
| mtstpLii138 | 244.18 | 92 | 22.46 | 9.5 |
| mtstpLii140 | 109.13 | 89 | 9.71 | 8.9 |
| mtstpLii141 | 73.66 | 94 | 6.92 | 9.3 |

#### 2.2 *O. elektroscirrha* transcript abundances

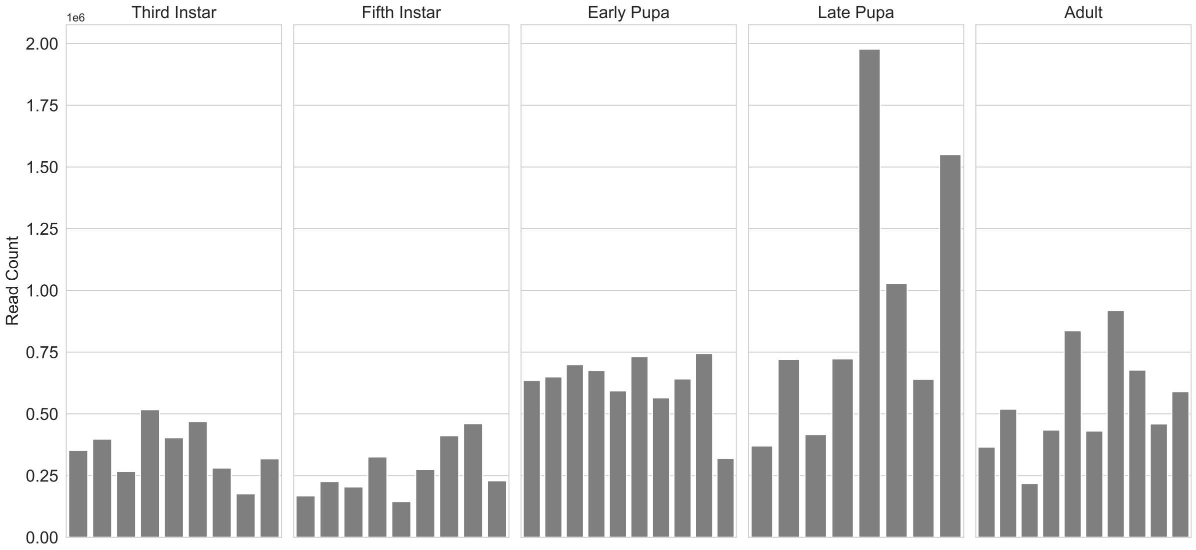

Figure S1: Quantifications of *O. elektroscirrha* read counts for each sample used in this study.

To see if our *O. elektroscirrha* transcript recovery was sufficient, we constructed a rarefaction curve for each sample to see where the number of genes we detected plateaued. This showed that even a sequencing depth as low as 140,000 sequences/sample allowed for sufficient gene detection. Therefore, this provides good evidence that our average of 538214.35 *O. elektroscirrha* sequences/sample was sufficient to describe their transcriptional activity.

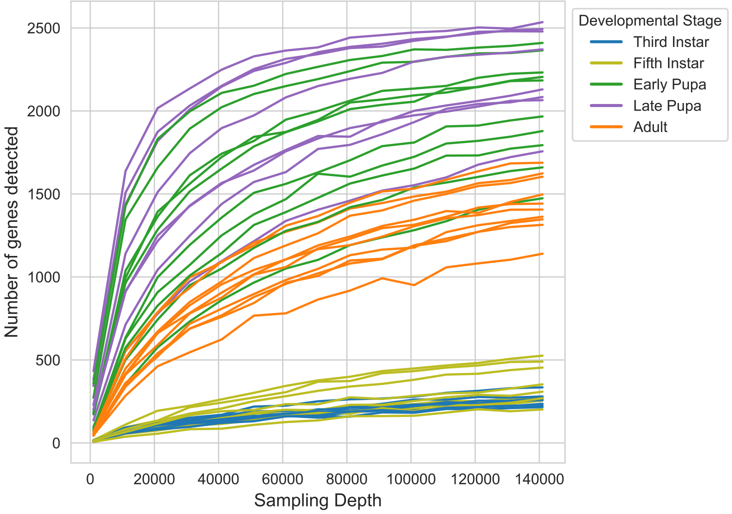

Figure S2: Rarefaction curve for *O. elektroscirrha* transcripts detected in each sample.

### 3 Supporting results

#### 3.1 The influences of host development and plant diet on *O. elektroscirrha* transcription

There was no significant difference in *O. elektroscirrha* transcription between *A. curassavica*- and *A. incarnata*-reared hosts (F = 0.9361, p = 0.379). There was a significant difference by developmental stage (F = 52.4847, p $\leq$ 0.001). Pairwise comparisons of adjacent stages showed *O. elektroscirrha* transcriptional dissimilarity between each *D. plexippus* stage.

Table S3: A table showing the results for each correlation between phylogenetic and expression pattern diversity that were summarized in the main text.

| *D. plexippus* Transition | F | p |
|---|---|---|
| Third instar larva - Fifth instar larva | 2.4650 | 0.005 |
| Fifth instar larva - Early pupa | 42.9714 | 0.001 |
| Early pupa - Late pupa | 89.5450 | 0.001 |
| Late pupa - Adult | 52.9031 | 0.001 |
